# Don’t throw out the sympatric species with the crater lake water: fine-scale investigation of introgression provides weak support for functional role of secondary gene flow in one of the clearest examples of sympatric speciation

**DOI:** 10.1101/217984

**Authors:** Emilie J. Richards, Jelmer W. Poelstra, Christopher H. Martin

**Affiliations:** Biology Department, University of North Carolina at Chapel Hill, Chapel Hill, North Carolina, United States of America; Biology Department, Duke University, Durham, North Carolina, United States of America

**Keywords:** adaptive radiation, introgression, gene flow, sympatric speciation

## Abstract

Genomic data has revealed complex histories of colonization and repeated gene flow previously unrecognized in some of the most celebrated examples of sympatric speciation and radiation. However, much of the evidence for secondary gene flow into these radiations comes from genome-wide tests, which tells us little about how gene flow potentially influenced sympatric diversification. Here we investigated whole genomes of Barombi Mbo crater lake cichlids for fine-scale patterns of introgression between species with neighboring riverine cichlid populations. We did find evidence of secondary gene flow into the radiation scattered across < 0.24% of the genome; however, the functional and genetic diversity in these regions paint no clear picture of how that variation could have contributed to the ecological and morphological diversity found in the lake. Our results suggest that either variation in novel genetic pathways introduced during secondary gene flow contributed to the radiation, or that secondary gene flow was predominantly neutral with respect to the diversification processes. We also found evidence for differential assortment of ancestral polymorphism found in riverine populations between sympatric sister species, suggesting the presence of a hybrid swarm in the past. While the history of gene flow and colonization appears to be more complicated than once thought, the lack of compelling evidence for secondary gene flow influencing diversification suggests that we should not yet rule out one of the most celebrated examples of sympatric speciation in nature.

## Introduction

Sympatric speciation, the extreme endpoint on the speciation-with-gene-flow continuum, is traditionally defined as the evolution of reproductive isolation without the aid of geographic barriers (Coyne and Orr 2004). Sympatric speciation has fascinated evolutionary biologists since Darwin for its illustration of the power of complex interactions between natural and sexual selection to create new species. Despite intense searches, very few case studies have been able to meet the rigorous criteria for demonstrating sympatric speciation in nature (Coyne and Orr 2004; Bolnick and Fitzpatrick 2007). Even in some of the more convincing examples that do meet these criteria, genomic data have revealed more complex evolutionary histories of multiple colonizations and repeated gene flow than previously thought (Papadopulos et al. 2011; The Heliconius Genome Consortium et al. 2012; Geiger et al. 2013; Alcaide et al. 2014; Igea et al. 2015; Malinsky et al. 2015; Martin et al. 2015a; Kautt et al. 2016).

However, much of the support for complicated histories involving repeated gene flow events into radiations comes from genome-wide tests for gene flow (e.g. (Lamichhaney et al. 2015; Martin et al. 2015a; Meier et al. 2017)). One prediction of models of speciation with gene flow is that divergence between incipient species should be heterogeneous across the genome (Turner et al. 2005; Harr 2006; Feder et al. 2012; Nosil and Feder 2012a,b). Indeed, high heterogeneity in genomic differentiation has been found across the genomes of many recent or incipient sister species (e.g. Jones et al. 2012; Martin et al. 2013; Poelstra et al. 2014; Soria-Carrasco et al. 2014; Malinsky et al. 2015; McGirr and Martin 2016), although other processes besides differential gene flow across the genome can produce similar heterogeneous patterns (Noor and Bennett 2009; Nachman and Payseur 2012; Cutter and Payseur 2013; Cruickshank and Hahn 2014; Guerrero and Hahn 2017; Ravinet et al. 2017). Only a handful of genes may directly contribute to the speciation process whereas the rest of the genome is porous to gene flow while reproductive isolation is incomplete (Wu 2001; Wu and Ting 2004). Therefore, gene flow detected at the genome-wide level from populations outside the sympatric radiation does not by itself constitute evidence that secondary gene flow was involved in the divergence process among incipient species and shaped the radiation.

The Cameroon crater lake cichlid radiations are some of the most compelling cases for sympatric speciation in the wild (Coyne and Orr 2004). The most speciose of these radiations is found in the isolated 2.3 km-wide volcanic crater lake Barombi Mbo (Trewavas et al. 1972; Schliewen et al. 1994; Schliewen and Klee 2004). Barombi Mbo hosts a radiation of 11 endemic cichlid species, many of which have clear morphological and ecological separation from other sympatric species (Schliewen et al. 1994). Some endemics have evolved unique specializations, such as the spongivore *Pungu maclareni* and deep-water hypoxia specialist *Konia dikume* (Trewavas et al. 1972). Other endemics, such as *Stomatepia mariae* and *S. pindu,* appear to be incipient or stalled species complexes with only slight morphological and ecological divergence at the extremes of a unimodal distribution of phenotypes (Martin 2012). However, evidence of differential introgression, weak support for Barombi Mbo monophyly, and differences in levels of shared ancestry with outgroup riverine populations from genome-wide RAD-seq data suggest additional secondary gene flow into the radiation after the initial colonization, casting doubt on one of the best examples of sympatric speciation in the wild (Martin et al. 2015a).

Here we dissect those signals of repeated gene flow to investigate their role in the radiation using whole-genome sequences. We performed exhaustive searches for all genetic patterns consistent with secondary gene flow into the ancestral Barombi Mbo population or into subclades after their initial divergence using machine learning to finely dissect phylogenetic signal across the genome and genomic scans to test for differential introgression. We find evidence of both shared introgression between sister species and across subclades in the radiation as well as differential introgression among sister species across small regions of the genome. However, functional and genetic diversity in these regions do not paint a clear picture of how introgressed variants may have contributed to speciation in these groups. Our results suggest that either 1) rare introgression of variants in novel genetic pathways contributed to the morphological and ecological diversity of the radiation (speciation with an allopatric phase), 2) secondary gene flow was predominantly or completely neutral and did not contribute to diversification in Barombi Mbo (sympatric speciation with gene flow), or 3) multiple colonizations of the lake before diversification brought in genetic variation that was then differentially sorted among incipient species (sympatric speciation from a hybrid swarm).

## Methods

### Sampling and Genome Sequencing

We sequenced whole genomes of 1-3 individuals from 10 out of the 11 species within the sympatric radiation of Oreochromini cichlids in Cameroon crater lake Barombi Mbo (excluding *Sarotherodon steinbachi* which is morphologically and ecologically similar to the other three *Sarotherodon* species), an endemic *Sarotherodon* species pair from Lake Ejagham, and outgroup *Sarotherodon* individuals from all three river drainages flanking the lake: Cross, Meme, and Mungo rivers (e.g. see map in (Schliewen et al. 1994)). Details on the collection, extraction, alignment to the *Oreochromis niloticus* reference genome, and variant calling protocols following the standard GATK pipeline are provided in the supplementary methods.

### Characterization of introgression patterns across the genome

First, we exhaustively searched the genomes for patterns of non-monophyletic Barombi Mbo relationships using the machine learning program SAGUARO (Zamani et al. 2013) to identify regions of the genome that contained relationships consistent with expectations from multiple colorizations and secondary gene flow into the radiation (i.e. paraphyletic/polyphyletic Barombi Mbo radiations). This method infers relationships among individuals in the form of genetic distance matrices and assigns segments across the genomes to different topologies without a priori hypotheses about these relationships. We partitioned the genome into a total of 75 unique topologies (well past the inflection point at 30 topologies where the percent of genome explained by each additional topology plateaus; Fig S1) to exhaustively search for relationships where subclades or individual Barombi Mbo species were more closely related to riverine populations than other species in the crater lake, suggesting sympatric speciation after a hybrid swarm (i.e. differential sorting of ancestral polymorphism) or secondary gene flow into this subclade (introgression). Details on the SAGUARO analysis and filtering strategies for calculating proportions are provided in the supplementary methods.

We also looked for evidence of differential introgression within subclades of the radiation on both a genome-wide and local level using *f_4_* statistics (Reich et al. 2009; Patterson et al. 2012; Pickrell and Pritchard 2012). The *f_4_* statistic tests if branches on a four-taxon tree lack residual genotypic covariance (as expected in the presence of incomplete lineage sorting and no introgression) by comparing allele frequencies among the three possible unrooted trees. We focused on tests of introgression with the two outgroup clades from our sample that came from two main clusters: riverine populations of *Sarotherodon galilaeus* in the Mungo and Meme rivers (MM) and riverine populations of *S. galilaeus* from the more distant Cross River (CR). Based on the tree ((P1, P2),(*S. galilaeus* MM, *S. galilaeus* CR)), *f_4_* statistics were calculated for combinations of species among a) *Stomatepia*, b) the *Konia + Pungu* subclade, and c) *Myaka myaka* with *S. linnelli* as a representative of its sister *Sarotherodon* group. This subset of groupings was chosen to make these analyses more tractable by focusing on species with unique trophic ecologies within the radiation. Genome-wide *f_4_* statistics were calculated using the fourpop function in Treemix (Pickrell and Pritchard 2012). Standard error was estimated by jackknifing in windows of 1,000 adjacent SNPs to account for linkage disequilibrium.

We characterized heterogeneity in introgression across the genome among these same combinations and investigated whether differential introgression contributed variation potentially important in the divergence between species by calculating *f_4_* statistics in 10-kb sliding windows. We did this with a modified version of the ABBABABA.py and genomics.py scripts that use population allele frequencies of biallelic SNPs (https://github.com/simonhmartin/genomics_general;(Martin et al. 2015b); our modified version is provided in the supplementary materials). Significance of *f_4_* values in sliding windows across the genome were evaluated using the 1% tails of a null distribution generated from permutations of the *f_4_* test. For more details on the sliding window calculations of *f_4_*, see supplementary methods.

For each of these regions, we looked for annotated genes using the well annotated NCBI *Oreochromis* Annotation Release 102 and searched their gene ontology in the phenotype database ‘Phenoscape’ (Mabee et al. 2012; Midford et al. 2013; Manda et al. 2015; Edmunds et al. 2016) and AmiGO2 (Balsa-Canto et al. 2016) for pertinent functions to the specializations and observed morphological differences among species, such as skeletal system or pigmentation.

### Directionality of introgression

The sign of *f_4_* does not indicate the directionality of introgression because of the lack of an explicit outgroup. For example, in the tree (P1,P2),(P3,P4)), a positive *f_4_* value indicates gene flow either between P1 and P3 or P2 and P4. We narrowed down the directionality of introgression detected in these regions using the *f*_*d*_ statistic, a modified version of the *D*-statistic that looks at allele frequencies fitting two allelic patterns referred to as ABBA and BABA based on the tree ((P1,P2),P3,O)), where O is an outgroup species in which no gene flow is thought to occur with the other populations (Martin et al. 2015b). Using two individuals of *Coptodon kottae* from another Cameroon crater lake as our distantly related outgroup population and the same riverine and Barombi Mbo population combinations described above, *f*_*d*_ values were calculated in 10-kb windows across the genome using the same script and window settings as in the *f_4_* tests of introgression. The variation in *f*_*d*_ values per region was higher than *f_4_* of similar window sizes, perhaps due to the high variance between windows in number of sites that fit the desired ABBA/BABA patterns given our outgroup, so we used only those *f*_*d*_ outliers (the top 2%) that overlapped with significant *f_4_* outliers as potentially introgressed regions from which we could narrow down the two populations in which gene flow occurred.

We also visualized the directionality of genome-wide introgression detected with the *f_4_* statistics using Treemix (v 1.13) (Pickrell and Pritchard 2012). Treemix estimates a maximum likelihood phylogeny of the focal populations and then fits a user-specified number of migration edges to the tree by comparing genetic covariances between populations. We ran Treemix with *S. galilaeus* as root, and with 0 through 20 migration edges. To determine the most likely number of migration events, we performed likelihood-ratio tests comparing each graph to that with one fewer migration event, starting with 1 versus 0 events, and took as the most likely value the first non-significant comparison.

### Comparison of patterns of introgression to patterns of genetic divergence and diversity

Reduced levels of genetic polymorphism in a population may indicate a strong selective sweep. We can look at introgressed regions found in only a single Barombi Mbo species for evidence that they have been adaptive, suggesting that secondary gene flow brought in variation potentially important for speciation. To examine genetic diversity in candidate introgressed regions, we calculated between-population nucleotide divergence (*D*_*xy*_) and within-population nucleotide diversity (p) for pairwise species comparisons among the Barombi Mbo focal species and the riverine outgroups. D_xy_ and p were calculated over the same 10-kb windows as the *f_4_* tests using the python script popGenWindows.py (https://github.com/simonhmartin/genomics_general; (Martin et al. 2015b); see supplementary methods for more details on these calculations).

## Results

### Widespread polyphyletic relationships in Barombi Mbo are scattered across small regions of the genome

After conservative filtering of segments to remove uninformative regions (see supplementary methods and Table S1), the Barombi Mbo cichlid radiation was a monophyletic group across 53% of the genome and only 0.6% was assigned to topologies indicating a polyphyletic Barombi Mbo. These polyphyletic relationships are consistent with many patterns, including secondary gene flow, incomplete lineage sorting, divergent selection, and ancestral population structure. The most prevalent topology spanned 38.2% of the genome and featured the expected species phylogeny for this group, in which all Barombi Mbo individuals form a single clade with distant relationships to outgroup riverine *Sarotherodon* populations in Cameroon (Fig. 1A). The second most prevalent topology (spanning 11.8% of the genome) featured identical evolutionary relationships, except for a much shorter branch leading to *S. galilaeus* Mungo and Meme River populations (Fig. 1B). Branch lengths produced by SAGUARO have no direct interpretation as an evolutionary distance (analogous to a neighbor-joining tree), but may be useful for comparison to similar topologies with different branch lengths, e.g. regions with higher divergence rates (Zamani et al. 2013).

**Fig 1.**
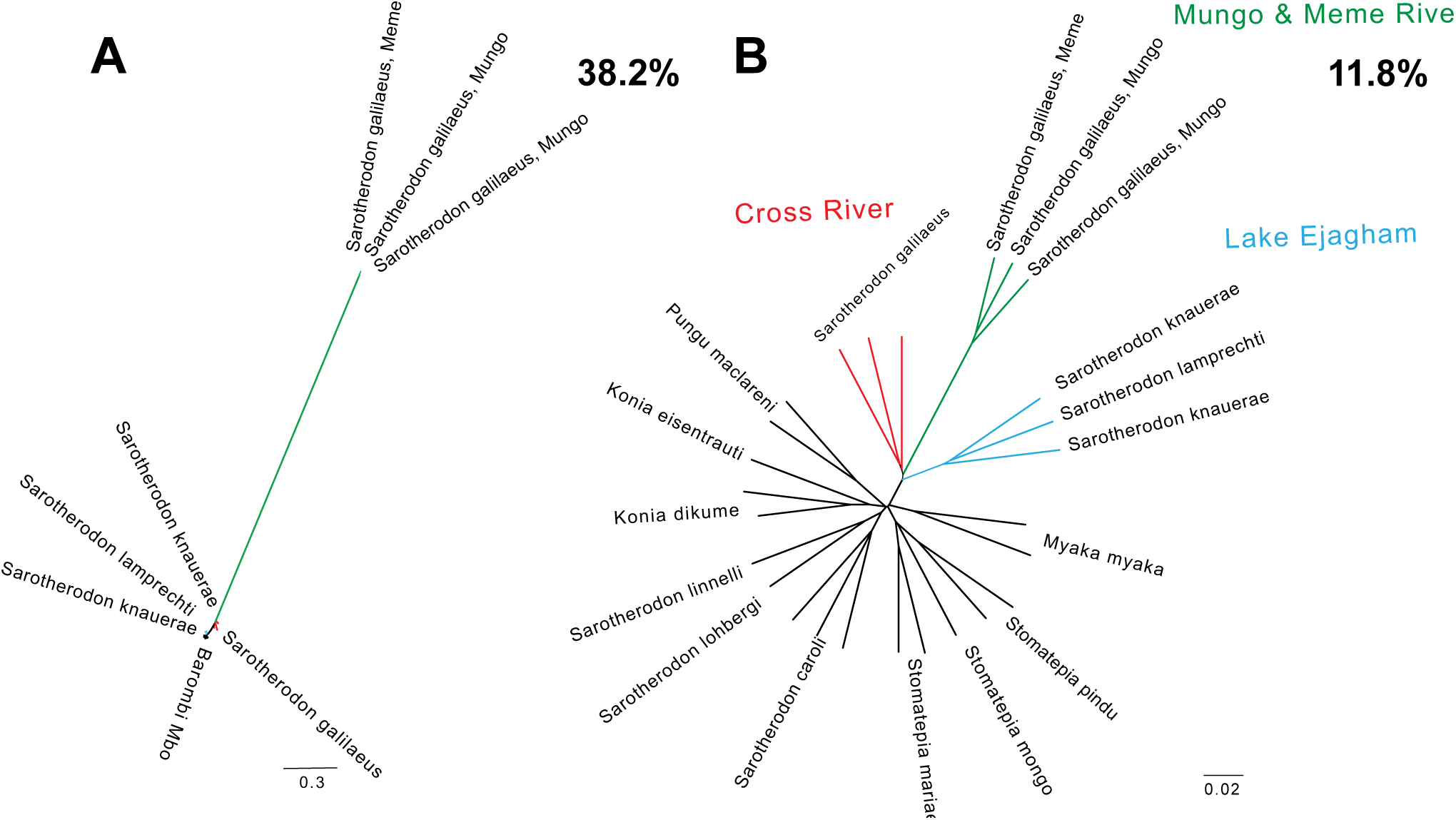
The most common topologies feature a monophyletic Barombi Mbo radiation. Across 96.2% of the genome Barombi Mbo species (black) are more closely related to each other than riverine outgroup populations of *S. galilaeus* Mungo and Meme River (green) and *S. galileaus* Cross River (red), or the Lake Ejagham *Sarotherodon* radiation (blue).

In 0.6% of the genome indicating polyphyletic Barombi Mbo relationships, we found evidence consistent with multiple colonizations of the lake. Since we were looking for patterns consistent with secondary gene flow or a hybrid swarm for subclades of the radiation, we focused on topologies where single species or entire subclades were more closely related to outgroups than other Barombi Mbo species, which represented only 0.24% of the genome. Some topologies featured an entire subclade (e.g. *Stomatepia*) as monophyletic, but more closely related to the riverine populations than other Barombi Mbo species, consistent with a hybrid swarm scenario before the diversification of the *Stomatepia* subclade. Other topologies featured individual species more closely related to outgroup riverine populations than sister species, consistent with secondary gene flow into that lineage after the initiation of divergence. For example, in *Stomatepia* we found topologies that group multiple species with riverine populations (Fig. 2A-B), but we also found topologies where individual *Stomatepia* species (*S. mariae* and *S. pindu*; Fig. 2C-D) were more closely related to riverine outgroups than other *Stomatepia*. In the *Konia* + *Pungu* subclade, we saw a similar pattern with topologies for the hypoxia and sponge-eating specialists (*K. dikume* and *P. maclareni,* respectively; Fig. 3A-B) but also a topology where the entire subclade was sister to the riverine outgroup populations (Fig. 3C). In the zooplanktivore *M. myaka*, we found topologies in which *M. myaka* was sister to the riverine populations (Fig. 4A-B), but also topologies where *M. myaka,* along with all the Barombi Mbo *Sarotherodon* species, were sister to the riverine outgroup populations (Fig. 4C-D).

**Fig 2.**
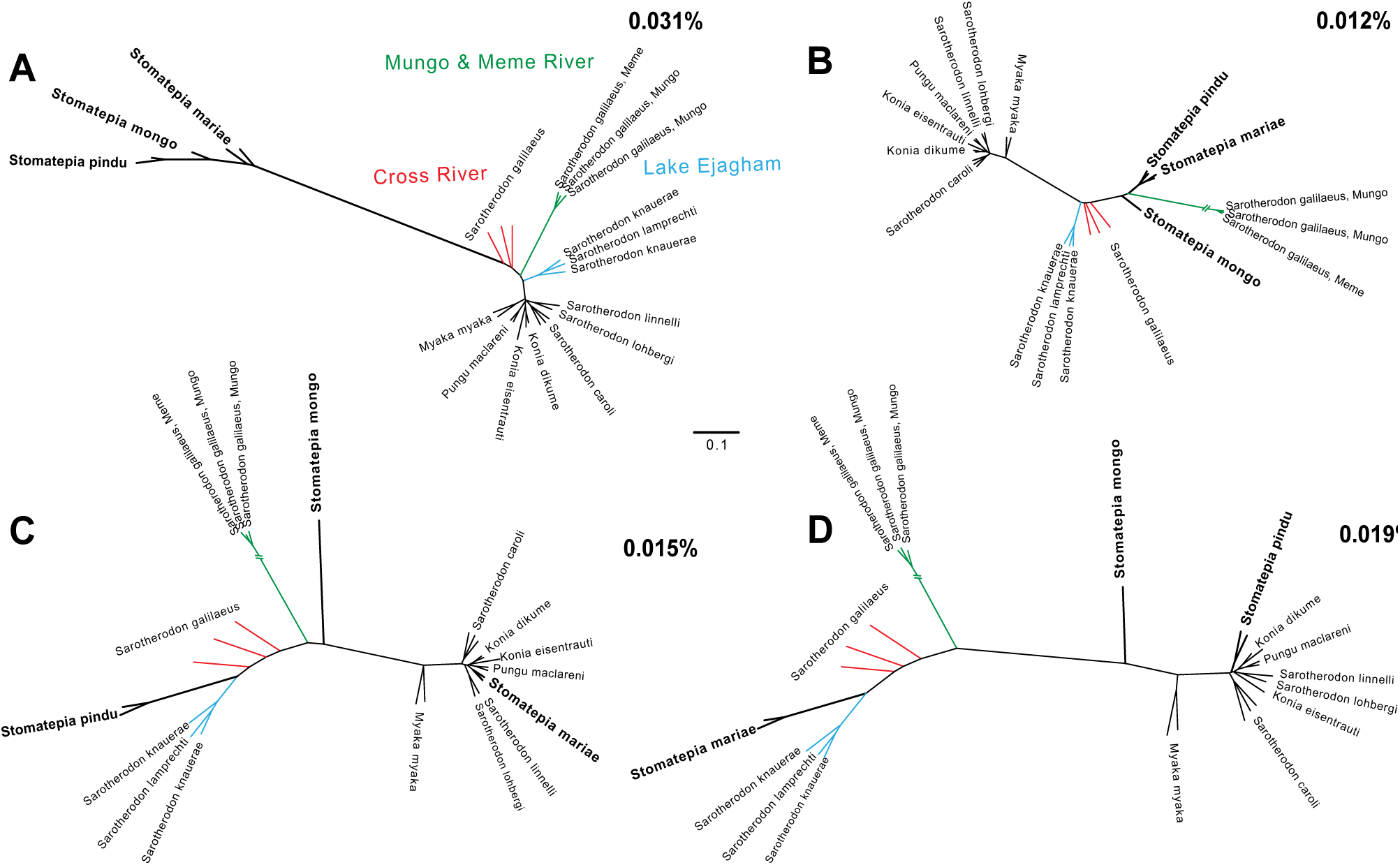
Topologies featuring Barombi Mbo polyphyly with riverine populations involving the *Stomatepia* three-species complex. Across small and independent proportions of the genome A-B) the entire *Stomatepia* clade, C) only *S. pindu,* and D) only *S. mariae* are more closely related to outgroups than other Barombi Mbo species.

**Fig 3.**
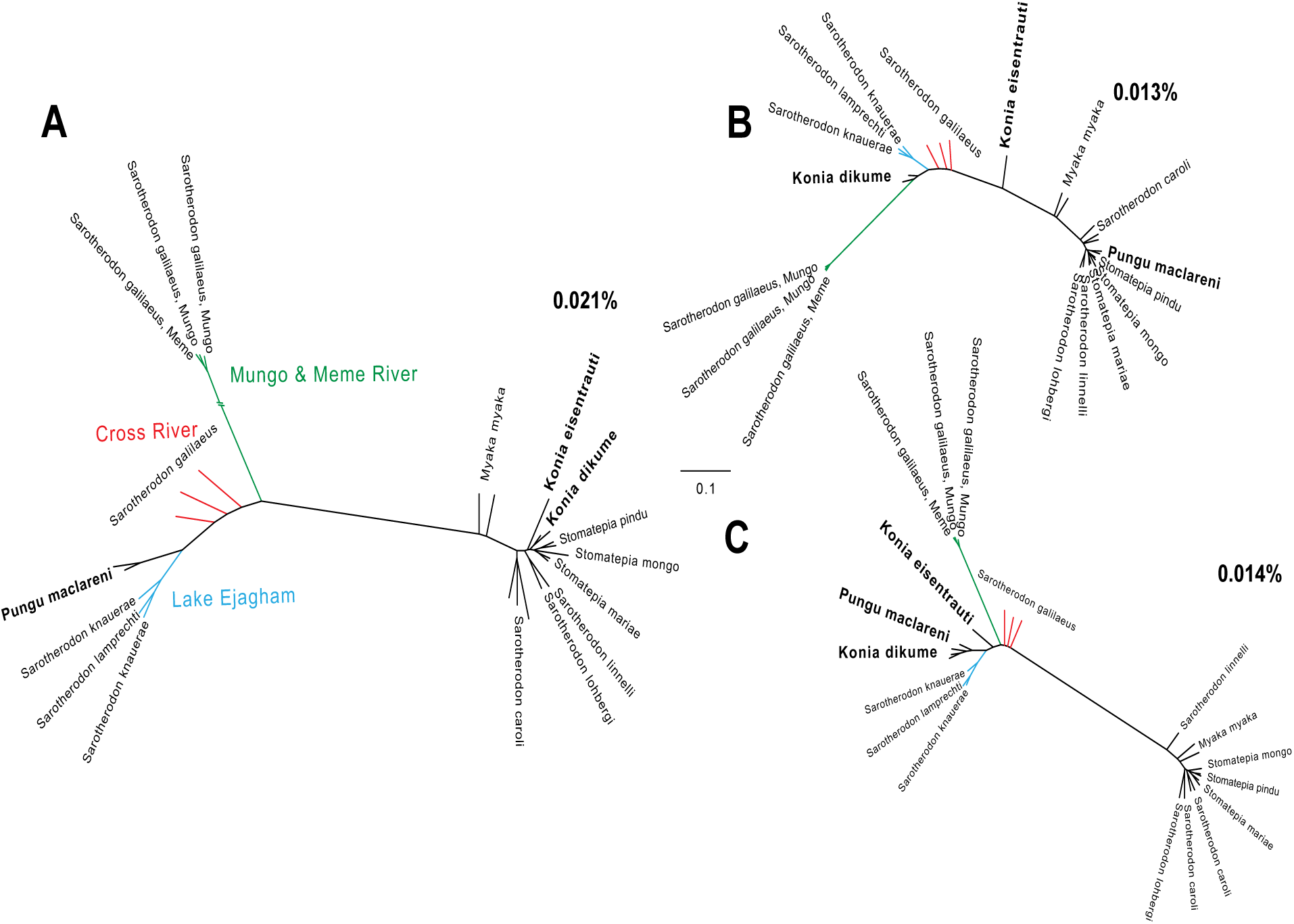
Topologies featuring Barombi Mbo polyphyly with riverine populations involving the *Konia + Pungu* subclade. Across small and independent proportions of the genome A) only *P. maclareni*, B) only *K. dikume*, and C) the entire *Konia + Pungu* subclade are more closely related to outgroups than other Barombi Mbo species.

**Fig 4.**
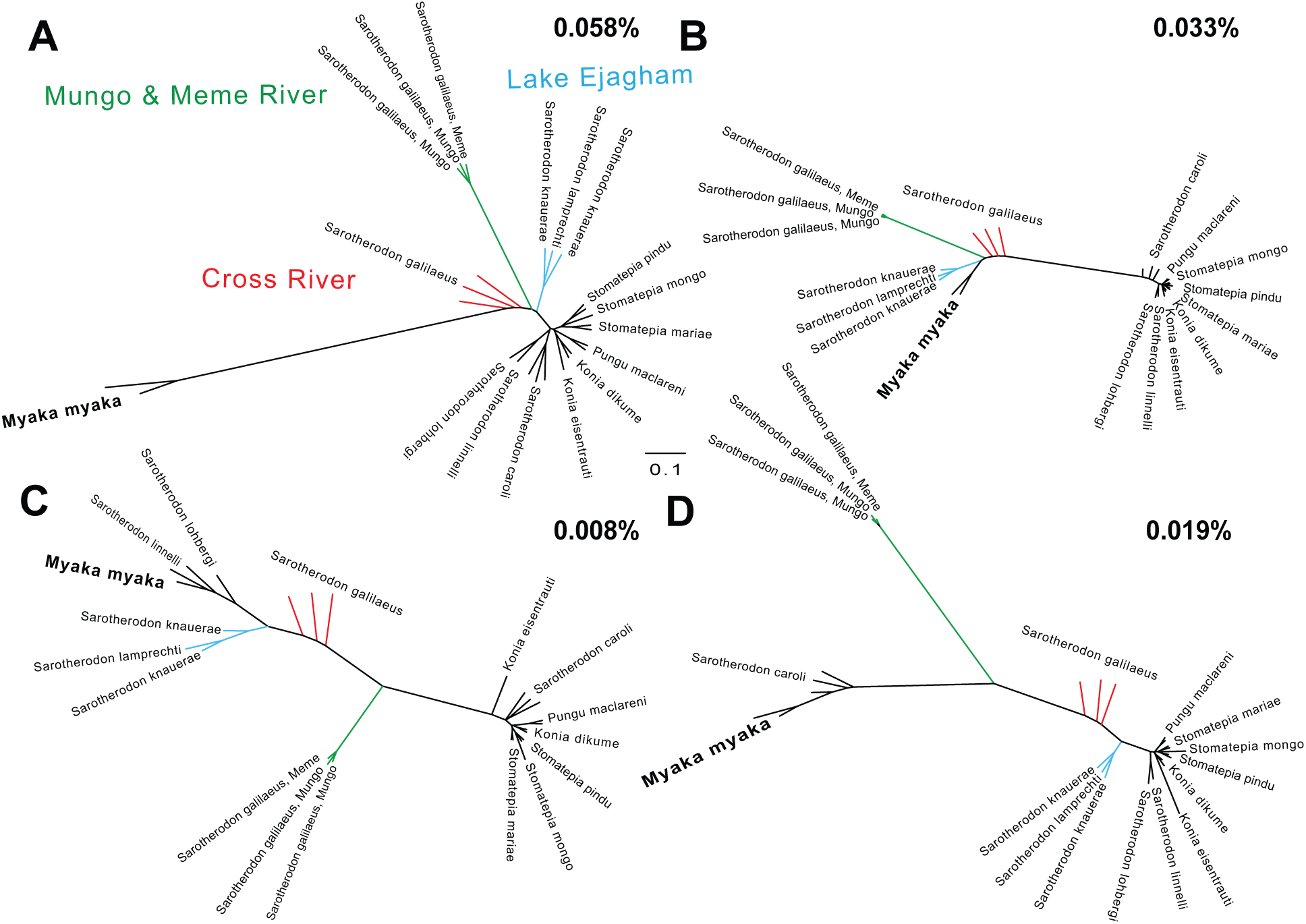
Topologies featuring Barombi Mbo polyphyly with riverine populations involving the *Myaka + Sarotherodon* subclade. Across small and independent proportions of the genome A-B) only *M. myaka*, C) *M. myaka* and two Barombi Mbo *Sarotherdon* species (*S. linelli* and *S. lohbergi*), and D) *M. myaka* and *S. caroli* are more closely related to outgroups than other Barombi Mbo species.

### Genome-wide evidence for differential introgression into the radiation

Consistent with evidence of differential introgression from RAD-seq data (Martin et al. 2015a), genome-wide *f_4_* tests provided evidence of genome-wide differential gene flow between some Barombi Mbo sister species and the outgroup riverine species (Table 1). There was significant evidence of genome-wide introgression in tests involving both *S. pindu* in the *Stomatepia* species complex and the hypoxia specialist *K. dikume* in the *Konia* + *Pungu* subclade. Some species pair combinations within these subclades did not show evidence of differential gene flow, suggesting that there may still be sympatric speciation occurring for some species, if not entire subclades. For example, there was no significant secondary gene flow detected genome-wide in the tests involving sister species *S. mariae* and *S. mongo* or *M. myaka* and *S. linnelli* (Table 1).

**Table 1.** Genome-wide *f_4_* statistics supporting differential introgression within Barombi Mbo radiation. Tests with significant evidence for differential introgression are highlighted in bold. The *f_4_* statistic was calculated for pairwise combinations among sister species of Barombi Mbo subclades and riverine populations of *S. galilaeus* from Mungo and Meme Rivers (MM) and Cross River (CR).

| Introgression with riverine outgroups:<br>(A,B) <-> ( MM, CR) | $f_4$ statistic | Z-score | P-value |
| --- | --- | --- | --- |
| *S. mariae*, *S. mongo* | $-2.04 \times 10^{-7} \pm -5.15 \times 10^{-7}$ | -0.39 | 0.69 |
| ***S. mariae*, *S. pindu*** | **$-1.92 \times 10^{-6} \pm -4.48 \times 10^{-7}$** | **-4.29** | **$1.8 \times 10^{-5}$** |
| ***S. mongo*, *S. pindu*** | **$-1.59 \times 10^{-6} \pm -4.98 \times 10^{-7}$** | **-3.19** | **0.0014** |
| *K. dikume*, *K. eisenrauti* | $-2.4 \times 10^{-6} \pm -6 \times 10^{-7}$ | **-4.01** | **$6.3 \times 10^{-5}$** |
| *K. eisenrauti*, *P. maclareni* | $-2.12 \times 10^{-7} \pm -6.15 \times 10^{-7}$ | 0.35 | 0.73 |
| ***K. dikume*, *P. maclareni*** | **$-2.56 \times 10^{-6} \pm -5.86 \times 10^{-7}$** | **-4.37** | **$1.2 \times 10^{-5}$** |
| *M. myaka*, *S. linnelli* | $-4.04 \times 10^{-7} \pm -7.11 \times 10^{-7}$ | 0.56 | 0.57 |

We also found evidence for widespread gene flow connecting populations across Barombi Mbo and neighboring riverine populations in highly interconnected population graphs; the likelihood of each graph did not plateau until reaching 10 admixture events (Fig S5). On the Treemix population graph with 10 admixture events, gene flow from the Mungo/Meme River populations of *S. galilaeus* occurred directly into individual species *S. mongo* and *K. eisentrauti* rather than the ancestral node of their respective subclades (Fig S6). The proportion of admixture inferred for these two events (0.1% into *S. mongo* and 0.4% into *K. eisentrauti*) was similar to the small proportions of the genome assigned to topologies consistent with secondary gene flow in the SAGUARO analyses. These admixture events pointing to the tips of the graphs suggest secondary gene flow events between nearby riverine populations and individual species within the radiation. In all population graphs allowing up to 21 migration events, any admixture from outgroup riverine populations appears to be coming from the Mungo and Meme rivers rather than the Cross River, consistent with the closer geographic proximity of the former drainages.

### Very few genomic regions contain signatures of differential introgression between sister species

Very few regions of the genome introgressed into single species from outgroup riverine populations (Fig 5A-C). In *Stomatepia*, only one region introgressed from Mungo/Meme Rivers into *S. pindu* and only three regions into *S. mariae*, respectively, suggesting secondary gene flow after initial diversification of *Stomatepia* (Table 2). Similarly, secondary introgression occurred into the *Konia* + *Pungu* subclade (Table 2). However, there was also evidence of shared introgression signals among sister species across all three subclades, where two subclade sister species shared introgressed regions from a riverine population. Only 0.000017- 0.0000354% of the genome appears introgressed into a single species of a Barombi Mbo subclade. This number is smaller than suggested in the SAGUARO analysis perhaps due to the conservative significance cut-offs and window size choice for *f_4_* statistic and that relationships observed in the polyphyletic topologies are consistent with other patterns besides introgression.

**Figure 5.**
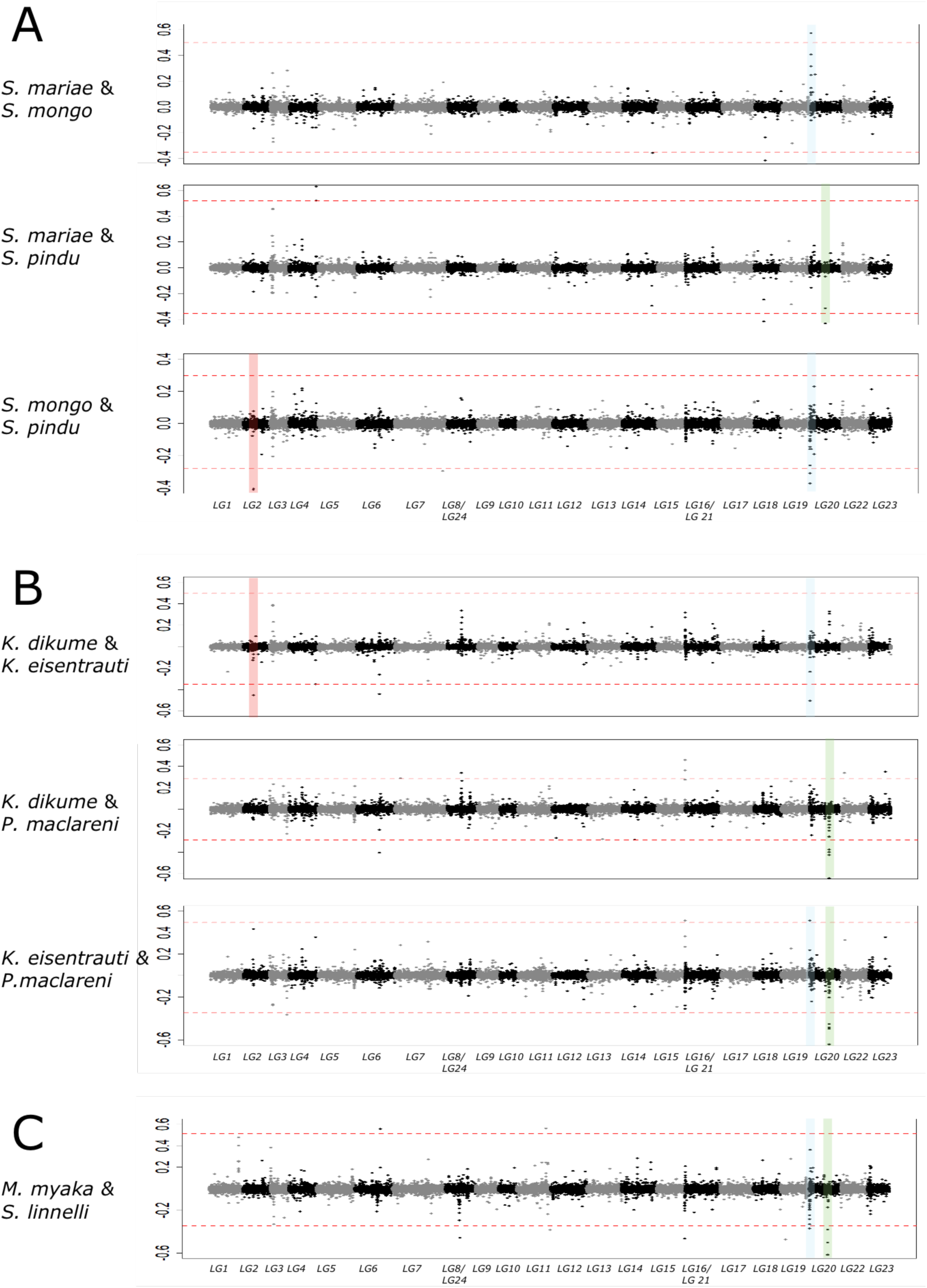
Manhattan plots of *f_4_* values between riverine populations of *S. galilaeus* from Mungo and Meme river and *S. galileaus* from Cross River and A) combinations of the three species of *Stomatepia*, B) combinations of the three species in the *Konia*+*Pungu* subclade, and C) *M. myaka* and a representative from its sister *Sarotherodon* clade: *S. linnelli.* Alternating gray/black colors indicate different linkage groups. Dotted red lines mark the permutation-based significance thresholds for each test (P = 0.02). Peaks highlighted in colors represent those signals of introgression shared across different subclades. Manhattan plots for the scaffolds not assigned to the 24 linkage groups are presented in Fig S2-4.

**Table 2.**
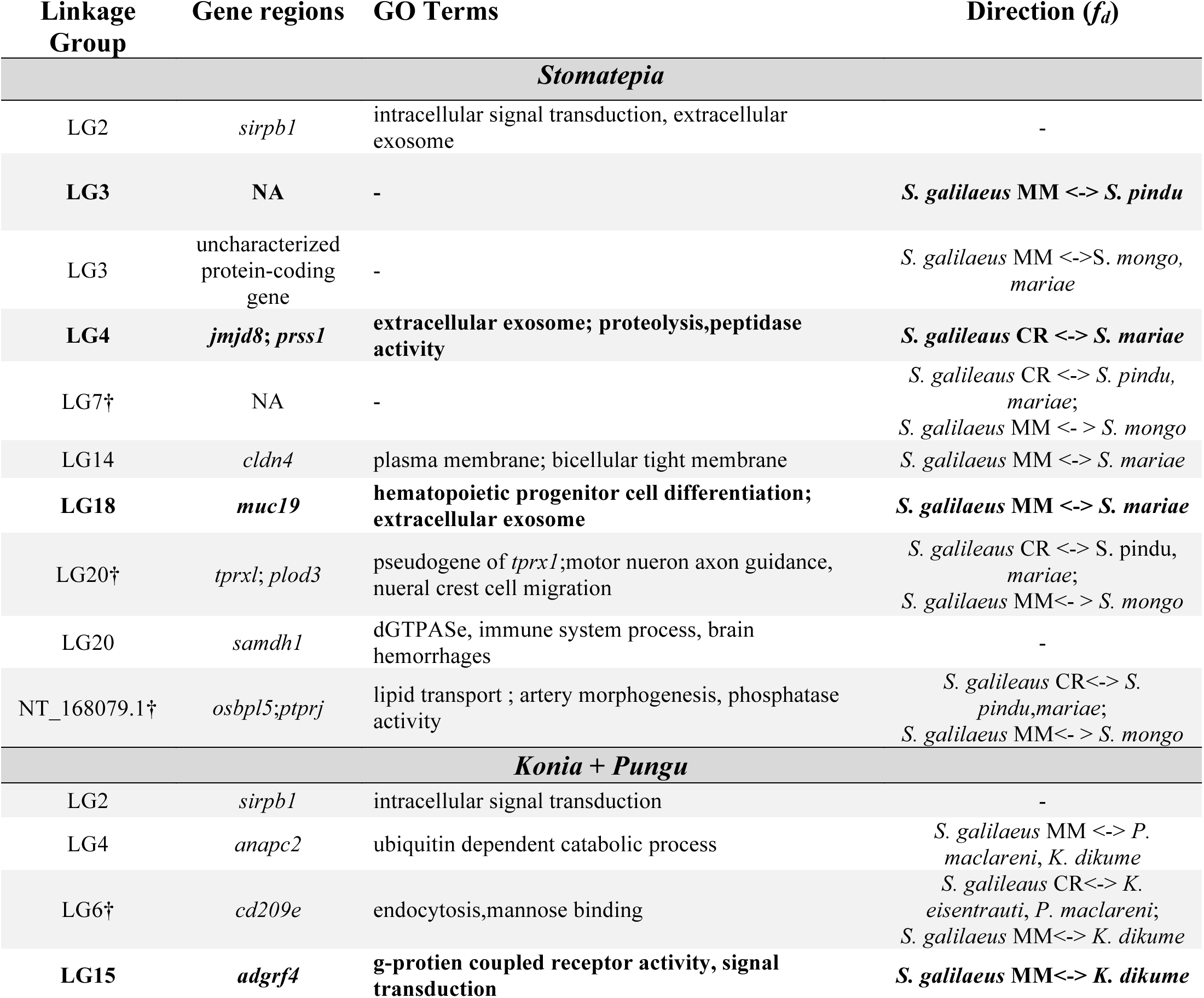
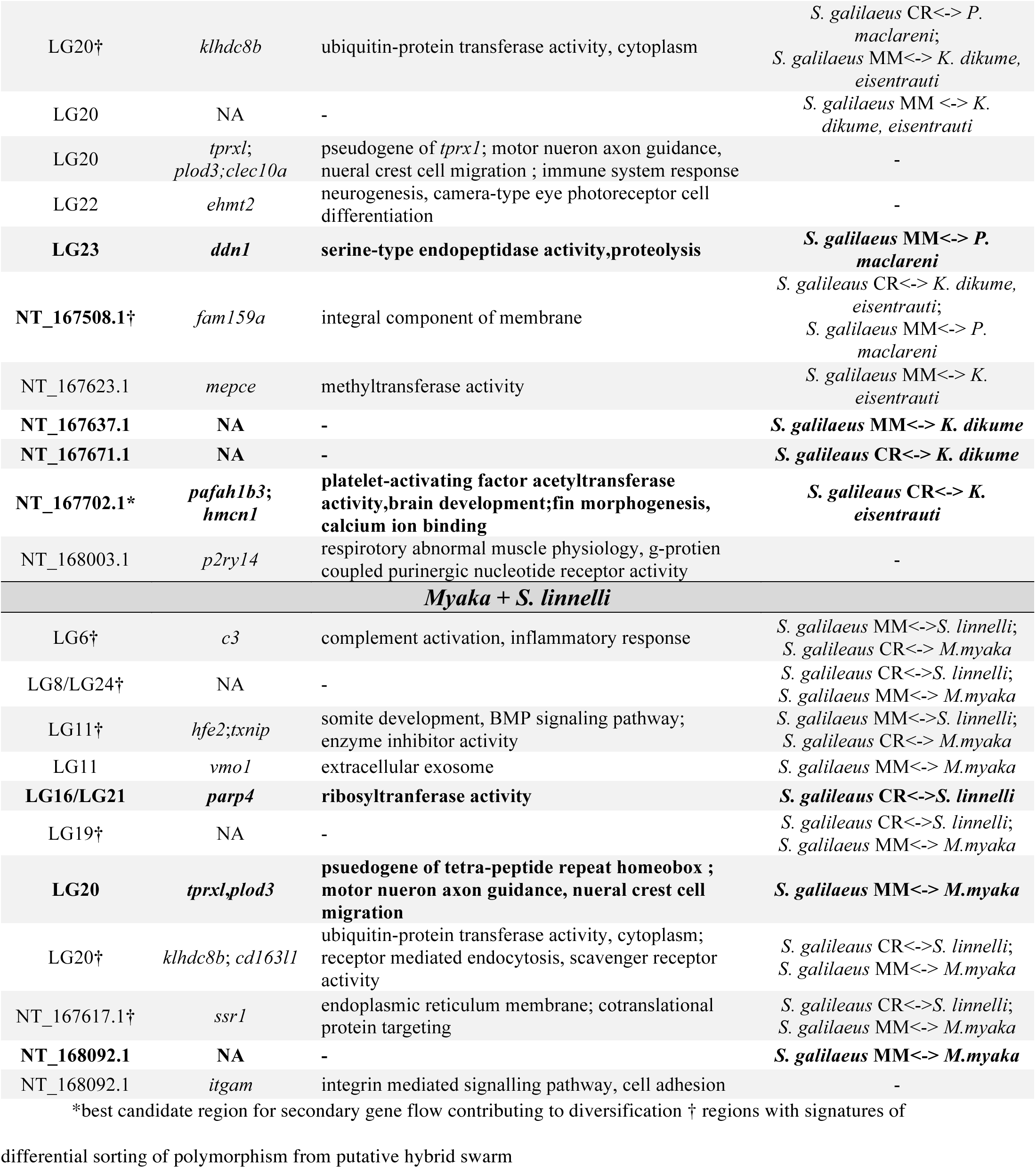
Candidate introgressed regions in Barombi Mbo cichlid radiation. These regions feature significant *f_4_* values between riverine populations of *S. galilaeus* (MM: Mungo and Meme River; CR: Cross River) and the three subclades of the radiation focused on in this study. Directionality was determined from *f*_*d*_ outlier values (above 98^th^ percentile) that overlapped with significant *f_4_* peaks. Unannotated regions, regions with no GO terms, and regions that were not *f*_*d*_ outliers are marked with (-). Regions introgressed into a single species within a subclade are highlighted in bold.

### Evidence for sympatric sorting of ancestral polymorphism within a hybrid swarm

A few of these 10-40 kb regions with peak signals of introgression were also present in multiple subclades, indicating differential assortment of introgressed variation shared among clades. For example, two significant *f_4_* outliers on linkage group 20 out of the 35 found across the genome appear within the *Stomatepia*, *Konia*, and *Pungu,* suggesting that some of this introgression may have occurred in the ancestral stages of the radiation and differentially sorted among species.

We also found 11 regions across the sister species pairs in which one species was more similar to one outgroup riverine population while its sister species was more similar to the other riverine population (Table 2). This signal is consistent with a hybrid swarm scenario due to multiple colonizations by riverine populations before diversification of some of the sister species and the sorting of polymorphisms brought in by these populations among incipient Barombi Mbo species. For example, two regions that appear to be differentially sorted between *S. galilaeus* and *S. mariae* and *S. pindu* from *S. galilaeus* CR versus *S. mongo* from *S.galilaeus* MM. Similar patterns were found scattered across the genomes for *K. eisentrauti* and *K. dikume* versus *P. maclareni* and *M. myaka* versus *S. linelli*.

### Weak support for functional importance of introgressed regions for species diversification

Although we did find evidence of differential introgression among sister species scattered across a small proportion of the genome, the types of genes found in these regions painted no clear picture of how introgressed variation may have contributed to speciation (Table 2). For example, differential introgression in *Stomatepia* occurred in regions with genes involved in a large range of biological processes, including intracellular signal transduction, immune system response, and motor neuron axon development (Table 2), with no obvious links to the highly divergent morphological, ecological, or patterning traits observed between these species (Martin 2012) nor to those traits normally associated with adaptive radiation in cichlid fishes such as body shape, pharyngeal jaw morphology, retinal pigments, or male coloration (Kocher 2004; Barluenga et al. 2006; Wagner et al. 2012; Brawand et al. 2014; Malinsky et al. 2015; Meier et al. 2017). Similarly, in both the *Konia* + *Pungu* and *Myaka* + *Sarotherodon* subclades, introgressed regions were near genes involved in a large range of biological processes not clearly associated with adaptive ecological traits in these species, such as *K. dikume*’s hypoxia tolerance, *P. maclareni’s* spongivory, and *M. myaka’s* zooplanktivory. For example, while there appears to be differential introgression in *Konia* in a region containing *pafah1b3*, a gene involved in platelet activation activity, and *K. dikume’s* deep water specialization includes higher blood volume with higher concentrations of hemoglobin (Green et al. 1973), it is not obvious how introgressed variation in *pafah1b3* would have played a role in the evolution of these traits from studies of its function in model organisms, which includes spermatogenesis and sterility in mice (Prescott et al. 2000; Koizumi et al. 2003; Yan et al. 2003).

Similarly, the amount of genetic diversity in introgressed regions does not suggest strong divergent selection on introgressed genetic variation due to hard selective sweeps. In line with the presence of peaks in *f_4_* values in these regions, between-population diversity (*D*_*xy*_) was typically high between one of the species and its sister species (Fig. 6). However, within-population diversity across many of these regions was often greater or comparable to scaffold and genome-wide averages (Table S2-4), suggesting these regions may not have experienced hard selective sweeps that would support their role in adaptive divergence among species (Fig. 6). In summary, although we found evidence for differential secondary gene flow between sister species in the radiation, we did not find strong functional support from gene ontology terms nor signatures of selection that the introgressed alleles were important for sympatric species diversification.

**Figure 6.**
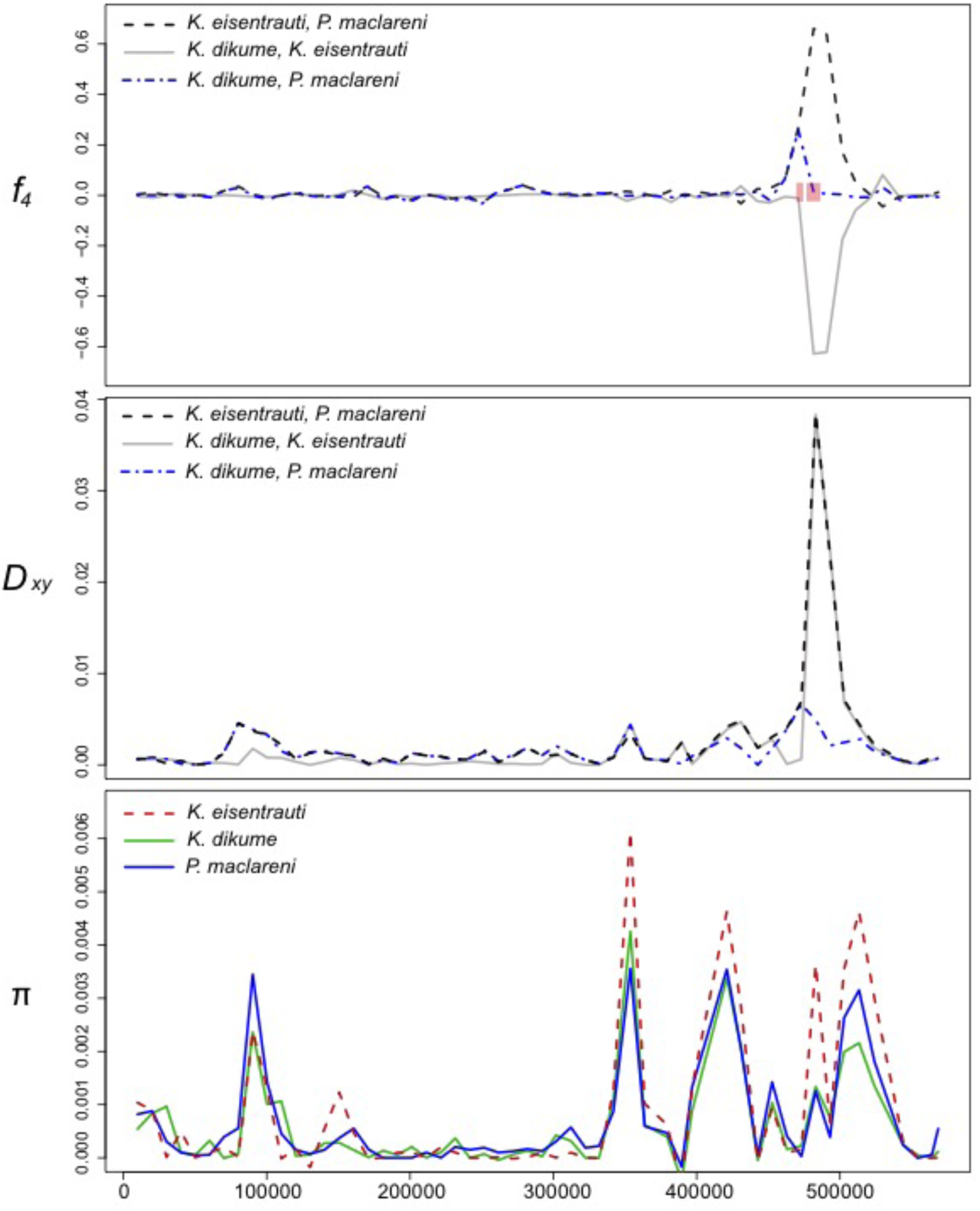
Candidate introgression region in the *Konia* + *Pungu* subclade of Barombi Mbo region containing genes *pafah1b3* and *hmcn1*. Row 1 shows the peak signal of introgression across scaffold NT_167702.1 detected from the *f*_4_ statistic across the three test combinations involving the three species in the *Konia* + *Pungu* subclade and riverine populations of *S. galilaeus* from Mungo, Meme River, and Cross River in non-overlapping 10-kb windows. The two genes in this peak are shown in red (*pafah1b3* on the left and *hmcn1* on the right). Row 2 shows between-population divergence (*D*_*xy*_) among the three combinations of sister species in the subclade calculated in non-overlapping 10-kb windows. Row 3 shows within-population diversity (*π*) in the same non-overlapping 10-kb windows.

## Discussion

### Little evidence that secondary gene flow promoted the diversification of Barombi Mbo cichlids

Our fine-scale investigations of introgression across the genomes of a celebrated putative example of sympatric speciation are consistent with two possible scenarios: 1) sympatric speciation in the presence of continuous neutral secondary gene flow into the radiation, or 2) speciation initiated by secondary gene flow. We found little support for the latter allopatric scenario from both a learning machine and sliding-window approach. From the SAGUARO analyses, our most conservative estimate of introgression into single species of the radiation ranges from 0.013 -0.019% of the genome. Estimates are similarly small from the *f_4_* statistics, ranging from 0.000017- 0.0000354% of the genome (Fig 7). Furthermore, even these significant outliers may represent false positives. First, our method of selecting introgressed regions from the 1% tails of a null distribution can always find outliers, even in the absence of introgression. Second, it is also difficult to distinguish signatures of differential introgression from the biased assortment of ancestral polymorphism into modern lineages, e.g. a hybrid swarm scenario that would still result in sympatric divergence entirely within the crater lake. Finally, even if our statistical outliers represent differentially introgressed regions, their importance to the speciation process is equivocal. We found no evidence of selective sweeps in these regions that would suggest they aided in divergence between species and they contain mainly housekeeping genes that do not clearly suggest how introgressed variation would have contributed to the radiation.

**Figure 7.**
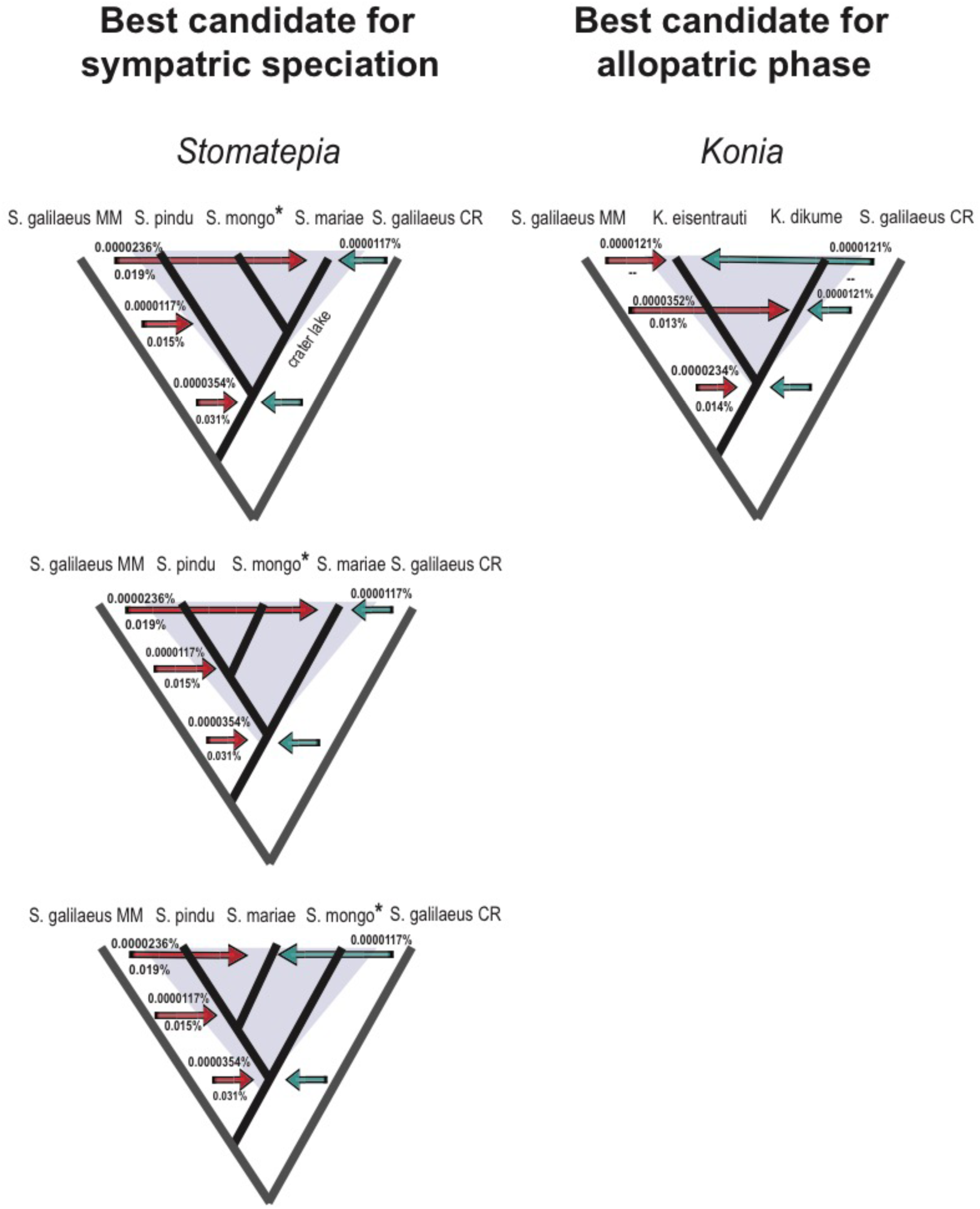
The contributions of differential and shared introgressed variation to the genomes of sister species in the genera *Stomatepia* and *Konia.* The left and right columns represent the best candidate sister species in the radiation for sympatric speciation and speciation with secondary gene flow respectively. Species with no evidence for differential introgression are highlighted with an asterick (*). The true sister species in the *Stomatepia* are unknown, so all three potential relationships among the complex are shown. Outgroup riverine *S. galilaeus* lineages from Mungo/Meme Rivers (MM) and Cross River (CR) are shown with gray branches. The percentage of introgression consistent with secondary gene flow into single species and a hybrid swarm from sliding window *f_4_* statistic tests is show by above arrows. Proportion of genome from two riverine sources (red: Mungo/Meme River; blue: Cross River) estimated by SAGUARO is shown below arrows. Introgression patterns not found in the SAGUARO analysis are represented with dashes (--). Blue arrows with no information that are aligned with red arrows represent patterns found consistent with hybrid swarm. Note that the separation of arrows along a branch is for the purpose of clarity and doesn’t represent known differences in the timing of introgression.

This contrasts studies on other systems using similar approaches which found compelling cases for adaptive introgression contributing to diversification (e.g. Abi-Rached et al. 2011; The Heliconius Genome Consortium et al. 2012; Huerta-Sánchez et al. 2014; Lamichhaney et al. 2015; Stankowski and Streisfeld 2015; Arnold et al. 2016; Meier et al. 2017), including our own previous work (Richards and Martin 2017). For example, several studies have found convincing candidate genes/variants in introgressed regions to suggest that adaptive introgression played a role in shaping ecological and morphological diversity. These include the detection of introgressed alleles linked to wing-color patterning involved in mimicry and mate selection in *Heliconius* butterflies (The Heliconius Genome Consortium et al. 2012), flower coloration involved in pollinator preferences for *Mimulus* species (Stankowski and Streisfeld 2015), and oral jaw size variation involved in scale-eating trophic specialization in *Cyprinodon* pupfishes (Richards and Martin 2017).

### Evidence for a hybrid swarm further complicates the role of gene flow in the speciation process in Barombi Mbo cichlids

Beyond speciation scenarios involving secondary gene flow, our findings also suggest another scenario for sympatric speciation in this system: sympatric speciation from a hybrid swarm involving the differential sorting of ancestral polymorphism among incipient species. A hybrid swarm is not easily detectable using the *f_4_* statistic because introgressed variation could be shared among diverging sister species, leading to an *f_4_* value of zero (Reich et al. 2009; Patterson et al. 2012). However, many of the *f_4_* peaks appear to be shared across at least two of the sister species in a subclade, shared between species of different subclades, or contain variation from both riverine populations (Mungo/Meme and Cross Rivers) that has been differentially sorted among sister species. All three of these patterns are consistent with an ancestral hybrid swarm before divergence between sister species occurred. This pattern of differential sorting of variation from a hybrid swarm from *f*_*d*_ analyses could also result from a lack of power in the statistic to distinguish the directionality of the introgression detected in those regions when using biallelic patterns and four populations (e.g. when two populations share similar allele patterns, the other two populations can share the opposite allele pattern by default). However, we also found evidence that entire subclades (e.g. *Stomatepia*) were more closely related to riverine populations than other Barombi Mbo subclades from the SAGUARO analyses that are also consistent with a hybrid swarm (e.g. Fig 2).

There are some caveats to our interpretations of secondary gene flow and its weak functional role in the ecological and morphological diversity observed within the lake. Recombination rate varies across genomes and determines the scale over which patterns of admixture and differentiation vary (Smukowski and Noor 2011). In our fixed sliding window size of 10-kb, we may have missed important patterns of introgression in regions of recombination hotspots, where such patterns are expected to be very fine-scale. Shared variation among species may reflect unsorted polymorphism from structured ancestral populations rather than hybridization. Introgression events can also be hard to distinguish from ongoing balancing selection of ancestral polymorphism that is sieved between species (Guerrero and Hahn 2017). While we focused on searching for genetic signatures of hard selective sweeps in introgressed regions, some of them with intermediate to high nucleotide diversity may have undergone soft selective sweeps, when selection drives multiple adaptive haplotypes to fixation. Some of these introgressed regions may have been adaptive and undergone soft selective sweeps, although the relative contributions of hard sweeps versus soft sweeps during adaptation and speciation is still the subject of much debate (Hermisson and Pennings 2005, 2017; Pritchard et al. 2010; Jensen 2014; Schrider et al. 2015).

### Best remaining cases for sympatric speciation within Barombi Mbo cichlid radiation

While the radiation as a whole may not have entirely arisen from sympatric speciation, some sister species within Barombi Mbo are better case studies of the process than others. Within the three-species *Stomatepia* subclade, there is little evidence that secondary gene flow played an important role in diversification. On a genome-wide level, we detected secondary gene flow in *f_4_* tests involving *S. pindu*. However, on a finer scale the one introgressed region unique to *S. pindu* is unannotated and the three introgressed regions unique to *S. mariae* contain four housekeeping genes involved in extracellular exosome activity and plasma membranes (*jmjd8*,*prss1*,*cldn4*,*muc19*). Shared signals of introgression among the three species represent a larger proportion of the genome than differentially introgressed regions, although both types of introgressed material appear to be rare in the genome (< 0.045%; Fig 7). The high ecological and morphological overlap among *Stomatepia* species suggests that this species complex may be stalled in the earliest stages of divergence. For example, *S. pindu* and *S. mariae* appear be at the extremes of a unimodal distribution of phenotypes; this is one major prediction of sympatric speciation models in the presence of only weak disruptive selection on ecological traits (e.g. (Matessi et al. 2002; Burger et al. 2006)), as measured in this species pair (Martin 2012).

Even for the two monotypic specialist species *M. myaka* and *P. maclareni*, there is minimal evidence for a role of secondary gene flow in the evolution of their trophic specializations. On a genome-wide level, the tests for differential introgression with one of the specialists and another species from the radiation were not significant. For the two regions which appear to be differentially introgressed between riverine populations and the zooplantivore *M. myaka,* one has no annotations, suggesting it may be neutral gene flow while the other is an introgressed region in other subclades, a signal of differential sorting of variation after a hybrid swarm. For spongivore specialist *Pungu maclareni*, we found only a single region differentially introgressed with riverine populations. This region contains *ddn1*, a gene involved in serine-type peptidase activity, proteins which are found ubiquitously in prokaryotes and eukaryotes.

Among all the ecologically divergent species pairs focused on in this study, *K. eisentrauti* and *K. dikume* are the least convincing as a putative case for sympatric speciation between sister species. Similar to *Stomatepia*, there is more evidence for shared introgression in regions of the *Konia* genome than differentially introgressed regions (Fig 8). However, differential introgression between the two *Konia* species occurs in regions with the best potential candidates in this study for contributing to diversification (although these regions are still not as strong as the ‘smoking guns’ observed in our past study of introgressed variation, e.g. Richards and Martin 2017). The gene *pafah1b3*, which is involved in platelet activation activity, is differentially introgressed between *Konia* species and may function in *K. dikume’s* deep water specialization of higher blood volume with higher concentrations of hemoglobin (Green et al. 1973). However, studies of the phenotypic effects of *pafah1b3* in model organisms do not hint at a role in hemoglobin concentration. Instead, mutations in this gene have been suggested to play a role in male spermatogenesis and fertility in mice (Koizumi et al. 2003; Yan et al. 2003). We also see differential introgression in a region containing the gene *ehmt2*, which is involved in neurogenesis and retinal cell differentiation. While it is not as directly clear how introgressed variation in this gene would have contributed to divergence among the *Konia* sister species, variation in visual traits such as color perception are important axes of diversification in other cichlid radiations, particularly along a depth gradient (Terai et al. 2002, 2006; Seehausen et al. 2008). These two species also exist in microallopatry; *K. eisentrauti* is an abundant detritivore along the shallow littoral region of the lake while *K. dikume* is a deep-water specialist on *Chaoborus* midge larvae which is only collected in deep-water gill nets (Trewavas et al. 1972; Schliewen 1994). Both species are mouthbrooders and likely breed in non-overlapping habitats.

### Conclusion

The complex history of colonization in the crater evidenced in this and a previous genome-wide study suggests allopatric phases of the speciation process in the radiation, which violates one of the strict criteria for demonstrating sympatric speciation in the wild (Coyne and Orr 2004). Nonetheless, from our fine-scale dissection of where in the genome these signals are coming from, we cannot point to a functional role for secondary gene flow in the speciation process across any of the subclades. This suggests that either variation in genes with undiscovered functional effects underlies the divergent ecologies and morphologies seen in the lake or that any secondary gene flow was neutral with regard to its role in the speciation process. We also find evidence to support sympatric speciation after a hybrid swarm that resulted from multiple colonizations of the lake, still consistent with a scenario of sympatric speciation through differential sorting of ancestral polymorphism. Disentangling the effects of a putative hybrid swarm from secondary contact events on the speciation process will require a better understanding of the timing of gene flow events compared to the diversification times of Barombi Mbo species. We found evidence for gene flow into the radiation both before and after initial diversification of subclades within the lake. Even without this information, weak support for a functional role of secondary gene flow in the radiation of Barombi Mbo cichlids suggests that we should not rule out this example of a sympatric radiation just yet.

## Acknowledgements

This study was funded by a National Geographic Society Young Explorer’s Grant, a Lewis and Clark Field Research grant from the American Philosophical Society, and the University of North Carolina at Chapel Hill to CHM. We gratefully acknowledge the Cameroonian government and the local chiefs of Barombi Mbo village and surrounding communities for permission to conduct this research. We thank Cyrille Dening Touokong and Jackson Waite-Himmelwright for field assistance and Nono Gonwuou for help obtaining permits.

## Author contributions

EJR wrote the manuscript. EJR conducted all analyses except for Treemix; JP conducted Treemix analyses. CHM collected samples and provided the genomic data. All authors contributed to the development of ideas presented in the study and revised the manuscript.

## Data Accessibility

All datasets used for this study will be deposited in Dryad and the NCBI Short Read Archive (SRA).

